# Collective computation and the emergence of hunter-gatherer small-worlds

**DOI:** 10.1101/2021.08.16.456574

**Authors:** Marcus J. Hamilton

## Abstract

Two defining features of human sociality are large brains with neurally-dense cerebral cortices and the propensity to form complex social networks with non-kin. Large brains and the social networks in which they are embedded facilitate flows of fitness-enhancing information at multiple scales, but are also energetically expensive. In this paper, we consider how flows of energy and information interact to shape the macroscopic features of hunter-gatherer socioecology. Collective computation is the processing of information by complex adaptive systems to generate inferences in order to solve adaptive problems. In hunter-gatherer societies the adaptive problem is how to maximize fitness by optimizing information processing given the energy constraints of complex environments. The solution is the emergent macroscopic structure of the socioecology. Here we show how computation is extended across social networks to form the decentralized knowledge systems characteristic of hunter-gatherer societies. Data show that hunter-gatherer bands of co-residing families constitute computationally powerful networks that are embedded within hierarchically modular social networks that form complex metapopulations bound by fission-fusion dynamics at multiple scales facilitating the flow of information far beyond local interactions. These dynamics lead to the emergence of hunter-gatherer small-worlds where highly clustered local interactions are embedded within much larger, but sparsely connected metapopulations. Hunter-gatherers optimize local energy budgets in small groups but maintain interactions with much larger social networks while avoiding many of the ecological costs.

## Introduction

In this paper we consider how computation, energy, and sociality interact to shape the macroscopic regularities of hunter-gatherer socioecology. Specifically, we focus on how the optimization of energy and information at the individual level scales up to constrain the large-scale organization and spatial structure of hunter-gatherer metapopulations. The processing of energy and information across levels of social organization is what we refer to here as *collective computation*.

More generally, collective computation is the ability of complex adaptive systems to compute solutions to problems by accumulating, aggregating, and deploying information across scales [11]. For example, the brain performs collective computations by aggregating the firing of individual neurons to perform complex behavioral responses to external stimuli [69, 25]. Collective computation plays a central role in Bayesian theories of the brain where sensory information the brain receives from interacting with the world is used to update models built from prior experience [28]. Inferences are then made by deploying updated information to generate increasingly accurate models of the world with obvious fitness consequences [30, 18, 20].

Computations are energetically expensive, and so the costs of information processing are integrated into the energy budgets of complex adaptive systems, from the increased metabolic costs of fueling large human brains [39] to the ecological costs of supporting populations [36]. These trade-offs lead to the optimizations we are interested in here. The costs and benefits of large brains play a central role in evolution of human ecology at all scales, from the scheduling of the human life history [48] and the optimization of foraging behaviors [49], to the formation of social networks and their distribution across landscapes [36, 37]. Indeed, the propensity of humans to form social networks has been central to the evolution of human ecology as cooperation and learning aggregates and amplifies the knowledge, skill, and experience accrued by individuals over their lifetimes.

Hunter-gatherer societies are perhaps definitive examples of complex adaptive systems [71]. Hunter-gatherer societies are collectives composed of multiple autonomous individuals seeking to maximize inclusive fitness by interacting with each other and their environments, which lead to the emergence of macroscopic regularities that appear in data as correlations across multiple levels of social organization and the environmental regulatory systems on which they rely [36]. Human behavioral ecologists have built deep mechanistic theories of these optimizations including foraging behavior, time allocation, social learning, parental investment, and patch residence time, many of which are summarized in [49]. While behavioral ecology models derive the optimizations, collective computation is the process by which statistically sufficient regularities are extracted from environmental signals and used to evaluate decision variables. The macroecological perspective we pursue here coarse-grains over these local optimizations to focus on how social groups solve the adaptive problem of predicting regularities in stochastic environments through collective computation.

A common feature of many ethnohistoric hunter-gatherer societies are *knowledge systems* [63] that integrate social, ecological, and environmental information into cohesive cultural belief traditions [3, 75, 9]. Here, knowledge accumulated over generations essential to survival and cultural identity are encoded into norms of behavior, craft, kinship, mythology, art, and ritual [65, 66, 76, 73]. The resulting traditions of belief are central to hunter-gatherer adaptations and are transmitted rigorously and systematically across generations. The “Dreaming” traditions common to many Aboriginal Australian hunter-gatherers provide a prime example [68]. In Dreaming cultures, from birth, individuals are embedded in living landscapes of mythical events and inherit custodial obligations to a particular region of the landscape and all the sacred knowledge it contains [21]. Landscapes are overlain by networks of songlines, or dreaming tracks, created by the epic journeys of Ancestral beings as they travelled across the country forming rivers, mountains, springs, plants, and animals [26]. Dreaming tracks often extend across the territories of neighboring groups and some traverse the entire continent. These songline networks form mental maps that link locations, people, and resources in space and can only be traversed by memorizing the appropriate song cycles [17, 64, 21] in a tradition often described as “singing up” the landscape [23]. Stars and constellations are then used to build star maps facilitating travel along dreaming tracks [59, 58, 31]. People thus have the ability to travel far beyond their familiar landscapes by learning the appropriate song cycles from the appropriate custodians of the landscapes they will be traversing. Dreaming tracks, star maps, and song cycles create multidimensional virtual worlds through which individuals navigate their social, biological, and physical environments. Thus, all individuals from birth are nodes in a vast network that extends across the entire continent, the properties of which are encoded in local belief systems and maintained for millennia [21]. These traditions allow individuals to build detailed inferential models of resources, people, and landscapes far beyond local experience. In this sense, a Dreaming tradition is a collective computation that solves the adaptive problem of detecting, extracting, and storing regularities from a dynamic, stochastic, fluctuating social and physical environment, by encoding accumulated information into culturally-inherited knowledge systems.

### Tinbergen’s four questions

The goal of this paper is to develop an intuition into the role collective computation and information flow plays in structuring hunter-gatherer socioecology at multiple spatial-temporal scales. To do this we focus on the cross-cultural analysis of hunter-gatherer societies and their comparison to non-human primates (hereafter primates, unless otherwise stated) and non-primate mammals (hereafter mammals, unless otherwise stated). A useful framework to address the ecological and evolutionary role of collective computation in hunter-gatherer societies is to consider Niko Tinbergen’s four questions [72]: 1) *Function*: what is the adaptive function of collective computation in hunter-gatherer societies? 2) *Causation*: how is collective computation performed in hunter-gatherer societies? 3) *Phylogeny*: what is the evolutionary history of collective computation in the human species? and 4) *Ontogeny*: how is collective computation integrated into the life history of hunter-gatherers?

In the remainder of this paper we examine collective computation in hunter-gatherer societies by quantifying how individual cognitive computation scales up in social groups to form what is sometimes termed the *collective brain* [56]. We begin by considering the computational scaling of mammalian, primate, and human brains. Next we derive a model to describe the allometric scaling of group size across mammals, primates, and hunter-gatherers. We then combine the scaling of individual computation and social groups to form collective brains and the resulting organization of hunter-gatherer socioecology. At the end of the paper we summarize the answers to Tinbergen’s questions.

## Data and methods

We use three comparative data sets of mammal brain composition, mammal species ecology, and cross-cultural hunter-gatherer socioecology. The first is data on mammal brain composition from Herculano-Houzel [41]. The second data set is a combined macroecological database of mammalian ecology compiled by the author from published sources. Mammal body mass, group size, population density, and home range size came primarily from the PanTHERIA database [47], with additional group size data from Jetz et al. [46] and home range data from Kelt and Van Vuren [50]. Primate group sizes came from Dunbar et al. [27]. Additional body mass data was extracted from the Amniote database [57]. Mammal brain mass data came from a combination of Isler and van Shaik [45], Sol et al. [67], and Barton and Capellini [2]. Data were examined and cleaned: obvious outliers and errors in the data sets were followed up through original sources, comparison to additional published sources and/or respected online sources, including Animal Diversity Web and the IUCN Red List of Threatened Species. The third data set is the Binford cross-cultural hunter-gatherer database [5], which includes social group size estimates at five levels of social organization, in addition to population density, territory size, and the net primary production of environments for 339 populations. As the sample sizes and scales of resolution differ widely across the three data sets we do not control attempt to control for phylogenetic autocorrelation in either the mammal or the hunter-gatherer data.

Statistical tests and figures are generated in the R statistical computing environment [70].

## Results

### Currencies, optimizations, and gambits

We focus on the fundamental currencies of energy and information and their optimization in hunter-gatherer populations. A common measure of energy in biological systems is metabolism, which is defined as the uptake, transformation, and expenditure of energy by an organism to fund the ecological demands of growth, maintenance, reproduction, and motility [10]. Collective computation is the natural informational counterpart to metabolism in complex adaptive systems. In this paper, we generalize the definition given by Brush et al. [11] where collective computation is the accumulation, aggregation, and deployment of information in order to make inferences about the world that solve adaptive problems.

The assumption, or phenotypic gambit, we make here is that fitness-maximizing foragers reduce the uncertainty in the models they construct of their worlds by updating prior beliefs with new information they extract from their environments. By definition, a net gain in information is a net decrease in model uncertainty (i.e., informational entropy) [69] leading to increased predictability and improved inferences of the world [30]. The goal is to minimize surprisal by maximizing the mutual information between the model used to generate inferences about the environment and the actual physical structure of the environment. Ethnographic examples may include models used to predict the location of resources in time and space, or macroscopic features of the environment used to inform mobility decisions, such as when to leave a patch. Thus uncertainty-minimizers optimize metabolic budgets by minimizing energy and time costs. By accumulating relevant information about the environment in the form of sufficient statistics, uncertainty-minimizers generate increasingly predictive models of their world, which are used to compute increasingly effective inferences. Larger groups of cooperators have the capacity to accumulate increasing amounts of information, but will necessarily incur increasing energy costs in finite environments. As such, group sizes, and the broader structure of social networks emerge from scale-dependent trade-offs between the benefits of information processing versus the ecological costs of maintaining the aggregate metabolic demand of group members.

### Computation and cognitive allometry

We begin by considering collective computation at the individual level. The fundamental units of computation in the brain are neurons, which transmit information to other nerve, muscle, and gland cells. Neurons are responsible for receiving and transmitting sensory input from the external world which is used to create and update models that allow organisms to make inferences of their environments [28]. The differential ability to compute accurate inferences about the world is central to biological fitness [69, 29].

While there are tight correlations between body mass, metabolic rate, and brain mass across mammals, humans have particularly large brains for an average adult body mass of 60 kg [54, 38]. However, the uniqueness of the human brain is the number of neurons in the cerebral cortex rather than the total number of neurons in the brain (∼ 86 billion) [39, 38, 1]. The number of neurons in the mammalian cortex correlates positively with cognitive ability measured as task performance in behavioral experiments [40]. Following Herculano-Houzel, here we use the number of cortical neurons in the mammal brain as a basic measure of cognitive ability, and thus cognitive computation. We use Herculano-Houzel’s data [41] to develop an intuition into the differential scaling of cognitive computation across mammals (denoted by subscript *m*), primates (denoted by subscript *p*), and humans.

We describe the scaling relationship between a dependent variable, *Y*, and an independent variable, *X*, as a power law, which has the mathematical form

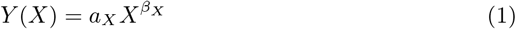

where *a*_*X*_ is a normalization constant and *β*_*X*_ is a scaling exponent describing how *Y* responds to a change in *X*; both parameters are time and scale invariant.

Figure 1A and Table 1 shows the scaling of brain mass, *B*, and body mass, *M*, where *B* ∝ *M* ^*β*^. For mammals *β*_*m*_ = 0.72 (0.67-0.77), and for primates *β*_*p*_ = 0.90 (0.74-1.06). Though primate brain mass increases with body mass faster than in other mammals, the difference between the slopes is not statistically significant at the 95% level (*t*_35_ = 1.97, *p* = 0.06).

**Table 1:**
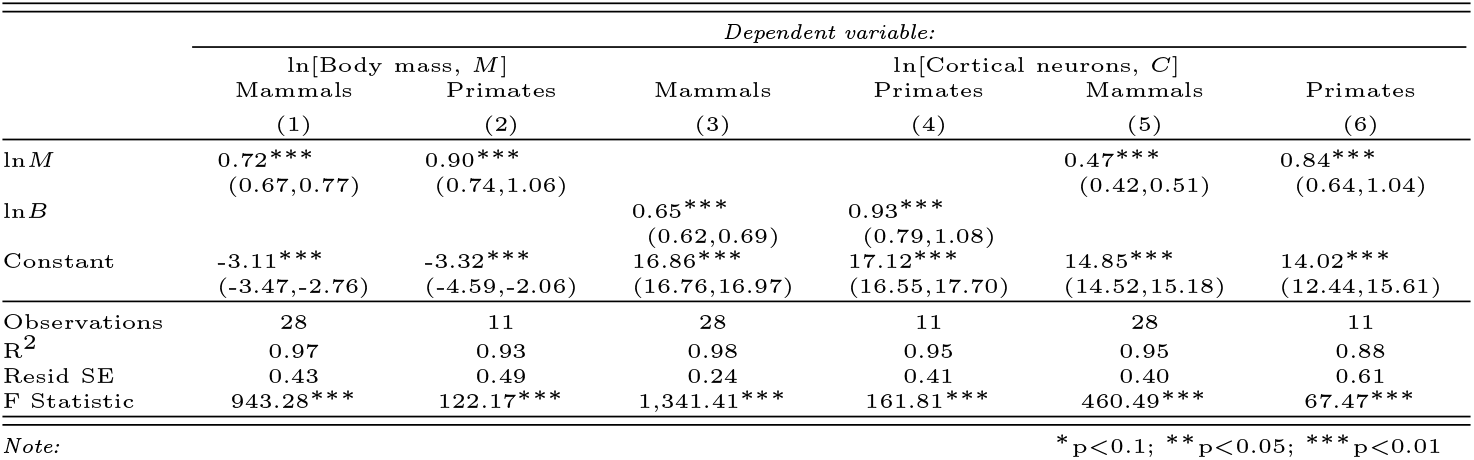
Brain mass, cortical mass, and cortical neuron allometry

**Figure 1:**
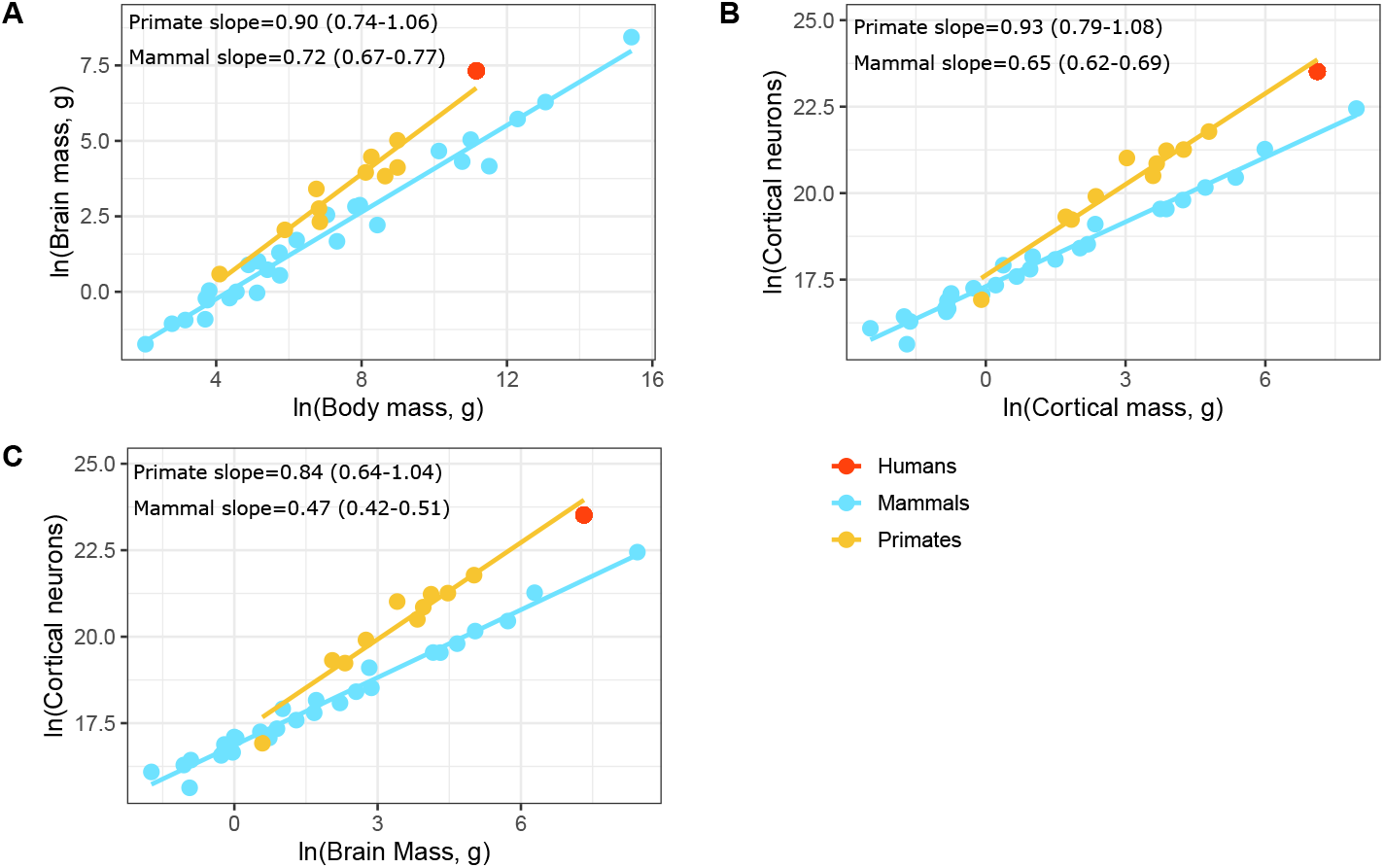
The allometric scaling of brain mass, cortical mass, and cortical neurons across non-primate mammals (blue points), non-human primates (orange points), and humans (red point) using data from Herculano-Houzel [41]. A) Brain mass increases with body mass slightly faster in primates than other mammals, though the difference in slopes is non-significant (see main text for statistical results). The human brain is large for an equivalently-sized mammal, but only slightly larger than expected for an equivalently-sized primate. B) The number of cortical neurons increases with cortical mass faster in primates than in other mammals, and so primate brains have a much higher density of neurons in their cortices than other mammals. C) The number of cortical neurons increases with brain mass nearly twice as fast in primates than in other mammals. Human brains have a near predictable number of cortical neurons for a primate, but more importantly they have more cortical neurons than any other mammal in the data set.

Figure 1B and Table 1 shows the scaling of the number of cortical neurons, *N* and cortical mass, *H*, where *N* ∝ *H* ^*α*^. For mammals *α*_*p*_ = 0.93 (0.79-1.08) and for primates *α*_*m*_ = 0.65 (0.62-0.69). Here, the number of cortical neurons increases with cortical mass significantly faster in primates than in other mammals (*t*_35_ = 1.97, *p* = 0.001), and so the density of neurons in primate cortices is significantly greater than in other mammals.

Figure 1C and Table 1 shows the number of cortical neurons, *N*, scales with brain mass, *B*, where *N* ∝ *B* ^*γ*^. For mammals *γ*_*p*_ = 0.84 (0.64 − 1.04) and for primates *γ*_*m*_ = 0.47 (0.42 − 0.51), so the number of neurons in primate brains increase significantly faster than in other mammals (*t*_35_ = 3.05, *p* = 0.004). Humans have the most cortical neurons of all species in the data set.

The number of cortical neurons in primate brains increases with body mass nearly twice as fast as other mammals, but it is important to note that humans have a predictable number of cortical neurons for a primate of 60 kg. Recalling Kleiber’s Law, *E* ∝ *M* ^3*/*4^, where *E* is the basal metabolic rate of a typical individual in a species, and *M* is the average adult body mass, then the cognitive return on whole organism metabolic investment in primates is 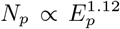, which is nearly twice that in other mammals, 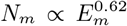. Moreover, the cognitive return on metabolic investment is superlinear in primates and sublinear in other mammals. As a consequence of Kleiber’s Law, larger body masses decrease reproductive rates but increase life spans [10] resulting in body-size invariant life-time reproductive effort across mammals [16] and humans [12]. However, Kleiber’s Law also describes economies of scale where mass-specific metabolic efficiency increases with body mass, and so natural selection will favor increased body masses if the result is to decrease mortality, even if reproductive rates are reduced. The brain mass scaling results presented here show that in primates larger body mass correlates with disproportionate increases in cognitive ability compared to other mammals. As a large-bodied primate, humans inherited an evolutionary legacy of neurally-dense cerebral cortices.

### Social group size allometry

To consider how cognitive ability scales up allometrically in groups, next we develop a macroecological model of group size. Social group size can be defined in many ways as there are many reasons social mammals live in groups [22, 53]. One method of estimating the average size of social groups is the average number of conspecifics (related or not) an individual encounters while performing daily tasks [47]. As such, we can consider group size, *g*, as a function of the encounter rate of an individual with conspecifics *λ* over the home range, *H*. Assuming the encounter rate of conspecifics, *λ*, is proportional to population density, *D*, we have

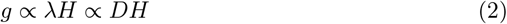

As is well-known in mammal ecology, the scaling of population density and body mass is described by Damuth’s Law [24], 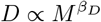 where *β*_*D*_ = −3*/*4, and home range scales as 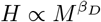, where *β*_*H*_ = 1 [46]. In Damuth’s Law population density is defined as *D* = *N/A*, where *N* is the number of individuals and *A* is a sampled area, measured in units *l*^2^, where *l* could be meters or kilometers. Note that we can equivalently write 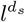, where *d*_*s*_ is the dimension of the area sampled. Therefore, if we consider a 3-dimensional environment, the area *A* in Damuth’s Law has an additional spatial dimension, and so can write 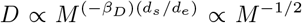, where *d*_*e*_ is the foraging dimension of the environment. Following equation 2 we then have an expression for group size as a function of body mass and the dimension of the foraging environment:

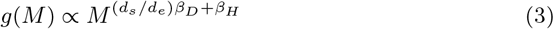

which in 2 dimensions gives *g* ∝ *M* ^1*/*4^, and in 3 dimensions *g* ∝ *M* ^1*/*2^. Equation 3 predicts that for a given body mass, mammal group sizes are larger in species that forage in three dimensions than species that forage in 2 dimensions, a prediction we confirm in data below.

We estimate these parameters from mammal species data. Figure 2A and Table 2 shows that for both primates and other mammals, home ranges scale approximately linearly with body mass, *H* ∝ *M* ^1^, and Figure 2B shows that population density in mammals scales as 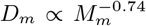, whereas for primates 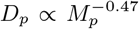 (see Table 2 for confidence intervals and test statistics). So, across primates and mammals home ranges scale approximately linearly with body mass, but within those home ranges, for a given body mass, population densities are higher in primates than other mammals, consistent with the dimension of their respective foraging niches. Figure 3A shows that, as predicted, primate group sizes scale with body mass as 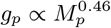, whereas social group sizes in other mammals scale a little shallower than expected, at 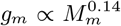. Interestingly, hunter-gatherer band sizes (the equivalent of mammal social group sizes in a home range), are considerably smaller than expected for a primate, but larger than expected for other mammals. Small band sizes would be consistent with the fact hunter-gatherers generally forage in niches that are close to 2-dimensional, with the possible exception of high canopy forests, fishing, or excavating roots, tubers, and fossorial animals, for example. As such, primates maintain larger group sizes by the increased resource supply rates and decreased competition of 3-dimensional foraging niches. Hunter-gatherer group sizes are constrained by 2-dimensional foraging niches leading to increased intraspecific competition exacerbated by the specialized food resources in the human diet. However, as we will explore below, hunter-gatherers integrate these small local groups into much larger networks facilitating flows of energy and information far beyond local interactions.

**Table 2:** Home range and population density

|  | *Dependent variable:* |  |  |  |
| --- | --- | --- | --- | --- |
| | ln[Home range, $H$] | | ln[Population density, $D$] | |
|  | Mammals | Primates | Mammals | Primates |
|  | (1) | (2) | (3) | (4) |
| $\ln M$ | 1.00***<br>(0.94,1.05) | 0.93***<br>(0.79,1.07) | -0.74***<br>(-0.79,-0.70) | -0.47***<br>(-0.60,-0.35) |
| Constant | -9.78***<br>(-10.18,-9.38) | -9.00***<br>(-10.13,-7.86) | 8.87***<br>(8.55,9.20) | 7.03***<br>(6.06,8.00) |
| Observations | 719 | 225 | 832 | 212 |
| $R^2$ | 0.65 | 0.42 | 0.55 | 0.22 |
| Residual Std. Error | 2.24 | 1.57 | 2.06 | 1.27 |
| F Statistic | 1,320.44*** | 164.72*** | 1,012.25*** | 58.39*** |
| *Note:* |  |  | *p<0.1; **p<0.05; ***p<0.01 |  |

**Figure 2:**
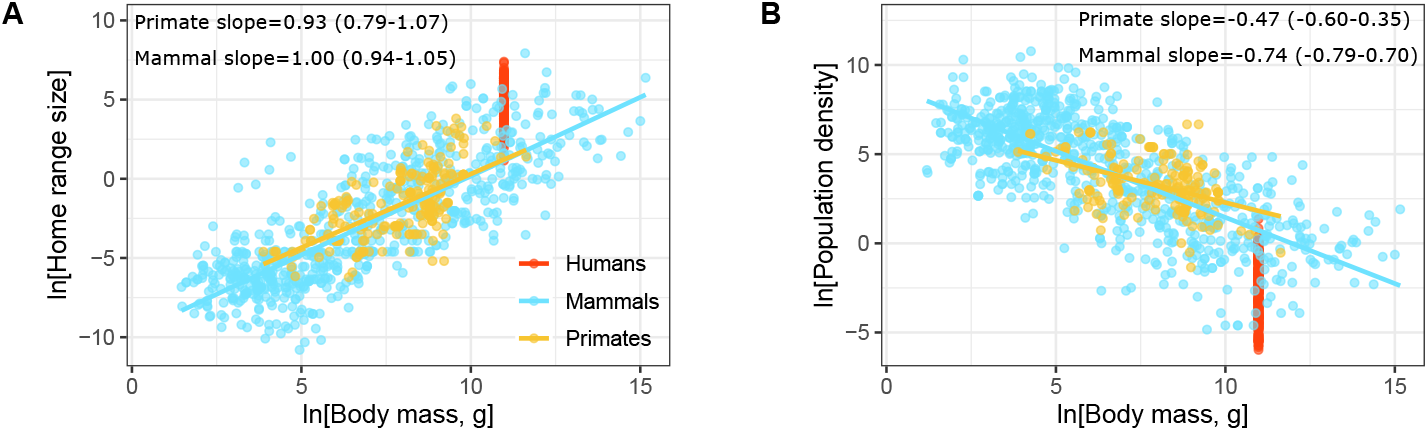
The allometric scaling of home range and population density in mammals, primates, and humans using data from PanTHERIA [47]. A) Home ranges increase approximately linearly with body mass in both primates and mammals (see Table 3 for details). Hunter-gatherers have large home ranges for their body mass. B) Primate population densities decrease with body mass slower than other mammals (i.e., Damuth’s Law), but hunter-gatherers have especially low population densities for both mammals and primates.

**Figure 3:**
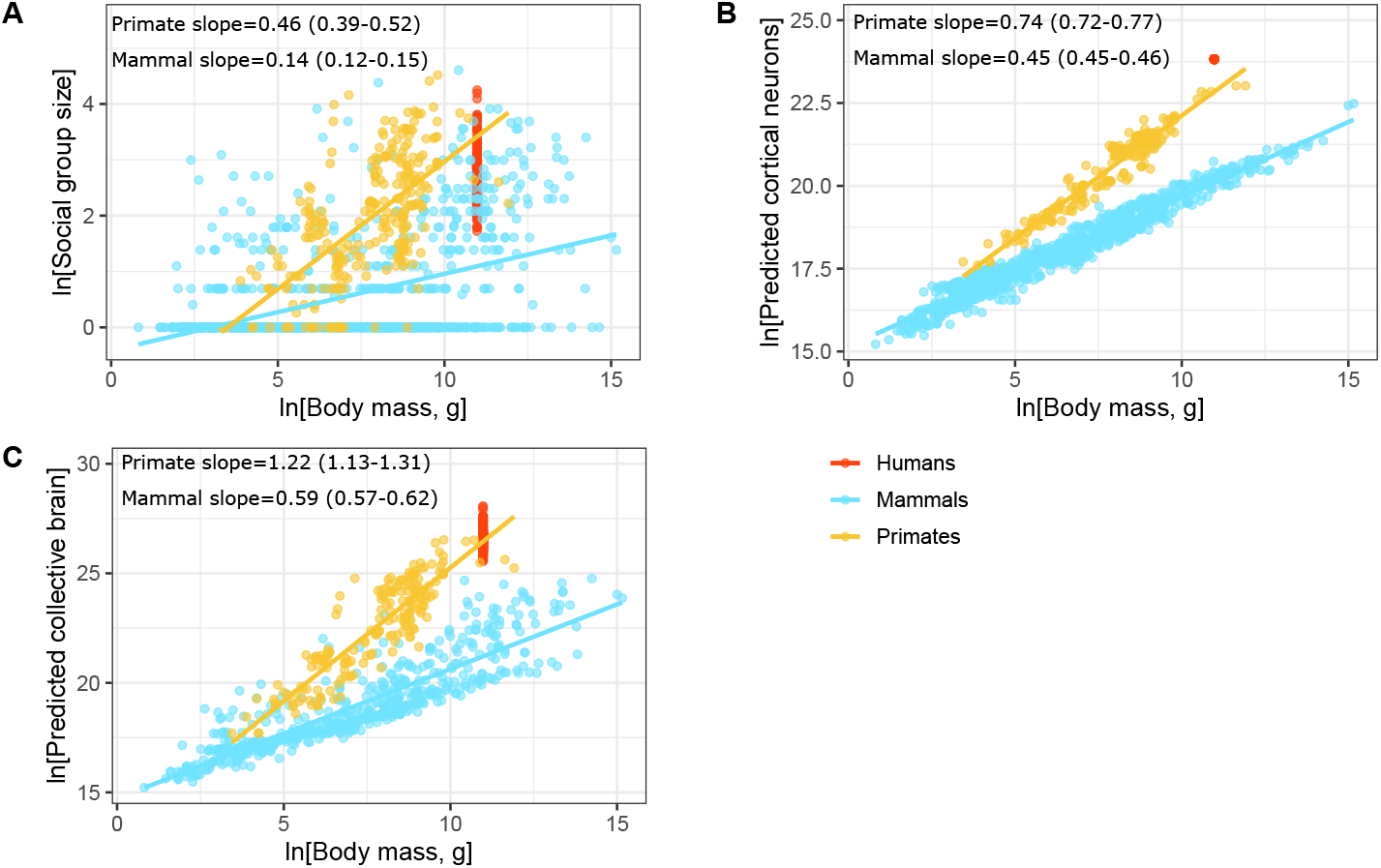
The allometric scaling of group size, (*g*), the estimated number of cortical neurons in the brain, (*N*), and the size of the estimated collective brain (*C* = *gN*) in mammals, primates, and humans. A) Social group size has a much stronger and faster (3-fold) response to body mass in primates (*β*_*P*_ = 0.46) than in other mammals (*β*_*M*_ = 0.14). Hunter-gatherer band sizes, which are the equivalent scale of social organization to the rest of the data, are considerably smaller than expected for a primate of equivalent body mass. B) Similar to Figure 1A the estimated number of cortical neurons increases with body mass slightly faster in primate brains than those of other mammals. C) Estimated collective brain mass increases sublinearly with body mass in mammals (*β*_*M*_ = 0.59) and superlinearly (*β*_*P*_ = 1.22) in primates. Hunter-gatherer collective brains are much as predicted for a primate despite having smaller than predicted group sizes and the largest of any mammal.

### Constructing the collective brain

Following Muthukrishna and Henrich [56] we use the term *collective brain* to refer to the collective computational power of a social group: the collective brain, *C*, is simply the product of the species-specific group size, *g*, and the average number of cortical neurons, *N*, in the brain of each individual in the group, and so *C* = *gN* . Following equation 3 we have a general expression for the collective brain:

**Table 3:** Group size, Estimated number of cortical neurons, and estimated collective brain mass

|  | *Dependent variable:* |  |  |  |  |  |
| --- | --- | --- | --- | --- | --- | --- |
| | ln[Group size, $g$] | | ln[Cortical neurons, $N$] | | ln[Collective brain, $gN$] | |
|  | Mammals<br>(1) | Primates<br>(2) | Mammals<br>(3) | Primates<br>(4) | Mammals<br>(5) | Primates<br>(6) |
| ln $M$ | 0.14***<br>(0.12,0.15) | 0.46***<br>(0.39,0.52) | 0.45***<br>(0.45,0.46) | 0.74***<br>(0.72,0.77) | 0.59***<br>(0.57,0.62) | 1.22***<br>(1.13,1.31) |
| Constant | -0.41***<br>(-0.54,-0.29) | -1.60***<br>(-2.11,-1.08) | 15.16***<br>(15.11,15.20) | 14.72***<br>(14.51,14.92) | 14.72***<br>(14.54,14.91) | 13.08***<br>(12.38,13.78) |
| Observations | 1,035 | 290 | 961 | 227 | 584 | 202 |
| R<sup>2</sup> | 0.20 | 0.39 | 0.96 | 0.94 | 0.79 | 0.79 |
| Resid SE | 0.83 | 0.85 | 0.27 | 0.30 | 0.92 | 0.99 |
| F Statistic | 265.02*** | 186.40*** | 22,038.22*** | 3,272.10*** | 2,210.72*** | 733.11*** |

Note:
\*p<0.1; \*\*p<0.05; \*\*\*p<0.01

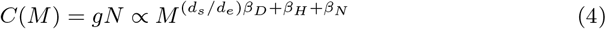

which in non-primate mammals predicts

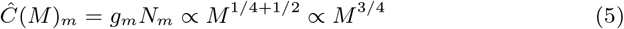

and in primates

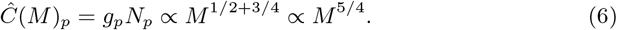

As such, collective brains in non-primate mammals are predicted to scale subinearly with body mass, whereas in primates collective brains are predicted to scale superlinearly. Given the scaling results in Tables 1 and 3 we find in mammals

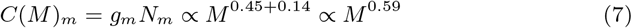

and in primates

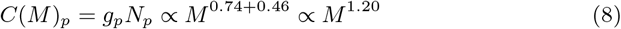

Empirically, the computational power of primate social groups increases with body mass twice as fast than in other mammals. To visualize this scaling relationship in Figure 3C we plot estimated collective brain mass (i.e., converting mammal brain masses in the data set to the estimated number of cortical neurons using the scaling parameters in Table 1) as a function of body mass in primates, mammals, and human hunter-gatherers. Hunter-gatherer bands have the predicted collective brain mass for a primate of 60 kg as small social group sizes (Figure 3B) are compensated by encephalized brains (Figure 1A). Figure 3C shows that hunter-gatherers have the largest collective brains of any mammal.

### Hunter-gatherer population structure

Next we examine the ways in which neighboring hunter-gatherer bands are connected and integrated into larger social networks. Hunter-gatherer populations are multi-scale societies where bands composed of co-residing families are connected to others across the landscape forming large-scale, low density, decentralized metapopulations. Empirically, these multiscale societies form self-similar, hierarchically modular networks [37] as shown in Figure 5. The kinship ties that bind families are extended to non-kin within co-residing bands by norms of reciprocal altruism and resource sharing [33]. Bands typically move multiple times over the course of a year as local foraging patches become depleted at rates predicted by environmental productivity [35]. Moreover, individuals move through the broader social network via several mechanisms: families may choose to relocate to another band or form a new band; individuals often visit friends and relatives in other bands; or husbands and wives change bands after marriage. As such, there is a constant demographic churn at the local level. At an aggregate level, in some environments (particularly hot and cold deserts) bands may seasonally disaggregate into individual families, and at other times local bands may aggregate into large temporary camps. Periodically, perhaps every few years, multiple bands may aggregate for mass events, such as ceremonies and rituals, or communal foraging events, such as rabbit drives or bison jumps (see [51, 13]). Hunter-gatherer metapopulations are connected by fission-fusion dynamics at all levels and timescales of the social network which serve to cycle information (both social and genetic) far beyond the daily interactions of individuals within bands. For example, the Ache foragers of northern Paraguay and the Hadza of Tanzania interact with on the order of 1,000 people over their lifetimes, which may constitute much of the metapopulation [43].

A hunter-gatherer metapopulation consists of Ω levels, which we label from the lowest level *ω*_1_ (families), the second level *ω*_2_ (bands), up to the highest level Ω, the metapopulation. The number of individuals, or average group size, at each level is denoted *g*_*ω*_, and so *g*_Ω_ is the total number of individuals in the metapopulation. Statistically, the branching ratio across all levels is constant, *θ* = *g*_*ω*+1_*/g*_*ω*_ ≈ 4, in which case hunter-gatherer metapopulations form self-similar hierarchically modular networks [37]: that is to say, on average we find four individuals in a family, *g*_1_, four families in a band, *g*_2_, four bands in a regional cluster of bands, *g*_3_, four clusters form a subpopulation, *g*_4_, and four subpopulations form the greater metapopulation, *g*_Ω_ (Figure 4A and Figure 5). As this branching ratio is an average across hundreds of populations, the branching ratio of any individual population may differ considerably from this overall pattern. Interestingly, however, the branching structure of these networks does not appear to vary across different environments. We see this invariance in Figure 4B and Table 3 where an ANOVA shows the average size of social groups at the five levels does not vary across ecosystem types, and in Figure 4C and Table 4 the regression model shows the variation within group sizes across the levels of the network is independent of local environmental productivity, measured as net primary production (g C m^-2^ yr^-1^).

**Table 4:** ANOVA table of social group size by organizational level and ecosystem

|  | Df | Sum Sq | Mean Sq | F value | Pr(>F) |
| --- | --- | --- | --- | --- | --- |
| Level | 4 | 3575.79 | 893.95 | 1361.89 | 0.0000 |
| Ecosystem | 6 | 45.28 | 7.55 | 11.50 | 0.0000 |
| Level:Ecosystem | 24 | 21.49 | 0.90 | 1.36 | 0.1129 |
| Residuals | 1156 | 758.80 | 0.66 |  |  |

**Figure 4:**
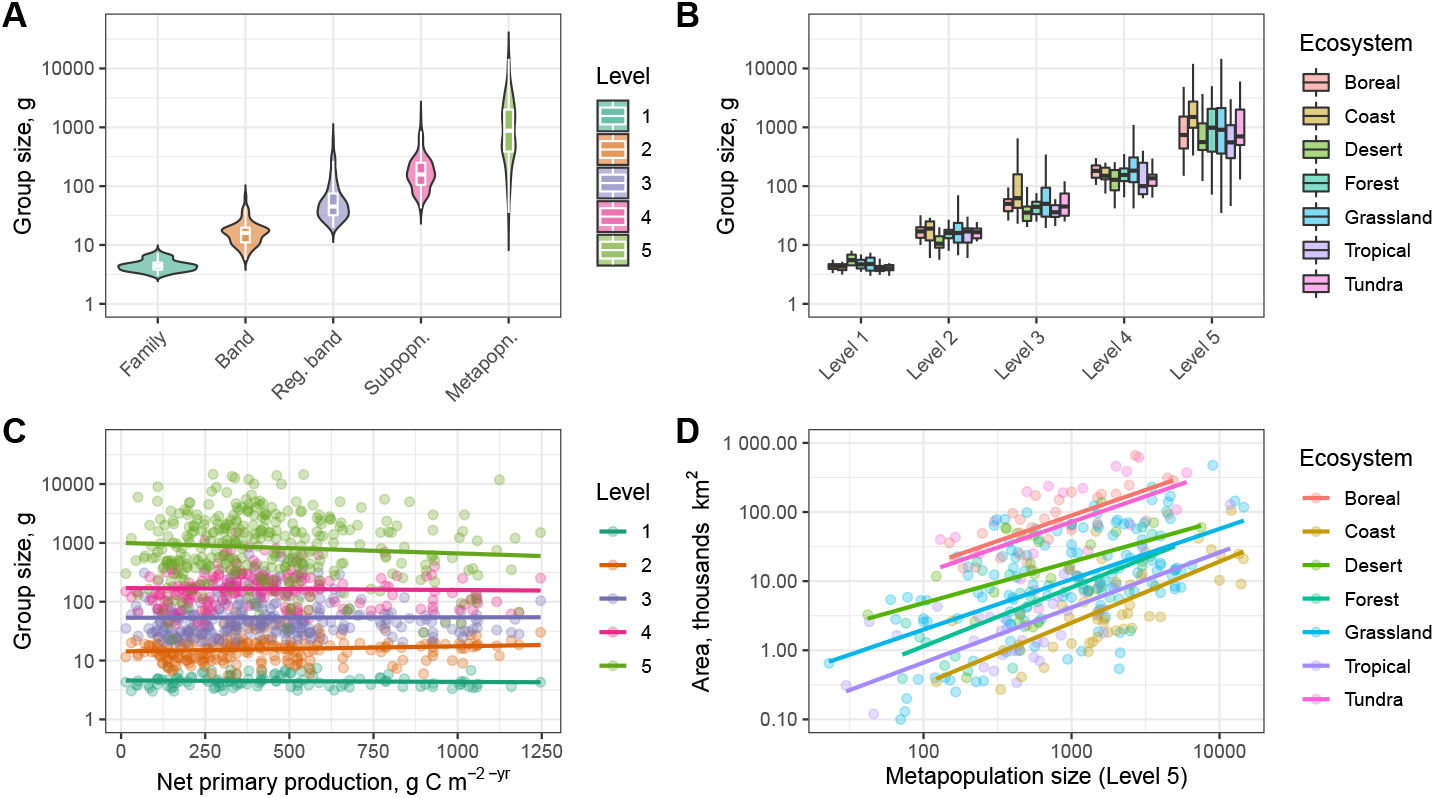
The hierarchically modular structure of hunter-gatherer networks using data from Binford (*n* = 339) [5]. A) Social group sizes at five levels of social organization for 339 populations shows a geometric series. B) The same social group size data at each of the five levels plotted by the ecosystem of the population, showing the geometric increase in group size holds across habitats. For statistical results, see Table 4. C) Social group size at each level plotted as a function of the net primary production of the environment for each population, showing that the variation in group size within each level is independent of environmental productivity. For statistical details see Table 5. D) Hunter-gatherer territory size plotted as a function of total population size (level 5), plotted by ecosystem type for each population. Across ecosystems hunter-gatherer populations are structured by self-similar networks, and exhibit economies of scale in space-use where larger networks. For statistical results, see Table 6.

Hierarchically modular networks have small-world-like properties with dense clusters of local connectivity and sparser global connectivity. Amongst individuals co-residing with multiple families in a band, daily interactions are likely frequent and ubiquitous, though structured by gender and age. If we assume all co-residing individuals, *g*, in a band, *ω* = 2, interact with each other, then the expected connectivity is given by the number of edges, *I*, which in a fully-connected bi-directional network is

**Table 5:** Hunter-gatherer social group size across levels by net primary productivity, *NPP*

|  | *Dependent variable:* |
| --- | --- |
| | $\ln[\text{Social group size, } g_{\omega}]$ |
| NPP | -0.0001(-0.001,0.0005) |
| Level 2 | 1.14*** (0.79, 1.48) |
| Level 3 | 2.45*** (2.12, 2.79) |
| Level 4 | 3.62*** (3.27, 3.97) |
| Level 5 | 5.38*** (5.06, 5.71) |
| NPP:Level 2 | 0.0003(-0.0004,0.001) |
| NPP:Level 3 | 0.0001(-0.001,0.001) |
| NPP:Level 4 | -0.0000(-0.001,0.001) |
| NPP:Level 5 | -0.0004(-0.001,0.0003) |
| Constant | 1.52*** (1.24, 1.80) |
| Observations | 1,191 |
| R<sup>2</sup> | 0.81 |
| Residual Std. Error | 0.83 |
| F Statistic | 572.35*** |
| *Note:* | * p<0.1; ** p<0.05; *** p<0.01 |

**Table 6:** Linear model of hunter-gatherer territory size by population size and ecosystem type.

|  | *Dependent variable:* |
| --- | --- |
|  | ln(Territory size, *A*) |
| ln*g*<sub>5</sub> | 0.74*** (0.32, 1.16) |
| ECOSYSTEM<sub>Coast</sub> | -4.59** (-8.19, -0.98) |
| ECOSYSTEM<sub>Desert</sub> | -0.56(-4.47, 3.35) |
| ECOSYSTEM<sub>Forest</sub> | -3.17* (-6.70, 0.36) |
| ECOSYSTEM<sub>Grassland</sub> | -2.28(-5.29, 0.72) |
| ECOSYSTEM<sub>Tropical</sub> | -3.46* (-6.92, -0.01) |
| ECOSYSTEM<sub>Tundra</sub> | -0.22(-4.63, 4.19) |
| ln*P*:ECOSYSTEM<sub>Coast</sub> | 0.15(-0.37, 0.66) |
| ln*P*:ECOSYSTEM<sub>Desert</sub> | -0.14(-0.73, 0.44) |
| ln*P*:ECOSYSTEM<sub>Forest</sub> | 0.11(-0.41, 0.63) |
| ln*P*:ECOSYSTEM<sub>Grassland</sub> | 0.02(-0.42, 0.47) |
| ln*P*:ECOSYSTEM<sub>Tropical</sub> | 0.06(-0.47, 0.58) |
| ln*P*:ECOSYSTEM<sub>Tundra</sub> | 0.001(-0.65, 0.65) |
| Constant | 6.29*** (3.47, 9.11) |
| Observations | 339 |
| R<sup>2</sup> | 0.58 |
| Residual Std. Error | 1.19 |
| F Statistic | 34.21*** |
| *Note:* | * *p*<0.1; ** *p*<0.05; *** *p*<0.01 |

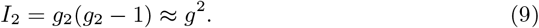

However, above the band level interactions between individuals in the metapopulation are far less frequent. If we assume minimal connections between bands then inter-band connectivity is proportional to the self-similar branching ratio *θ*, and so we have

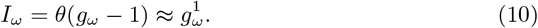

in which case inter-band connectivity is linear in *g*, for *ω >* 2. This leads to the structure shown in Figure 5, generated from the average properties of the Binford data used in this paper: internally bands are fully connected, but each band is minimally connected to other neighboring bands creating a statistically self-similar hierarchical modular network.

**Figure 5:**
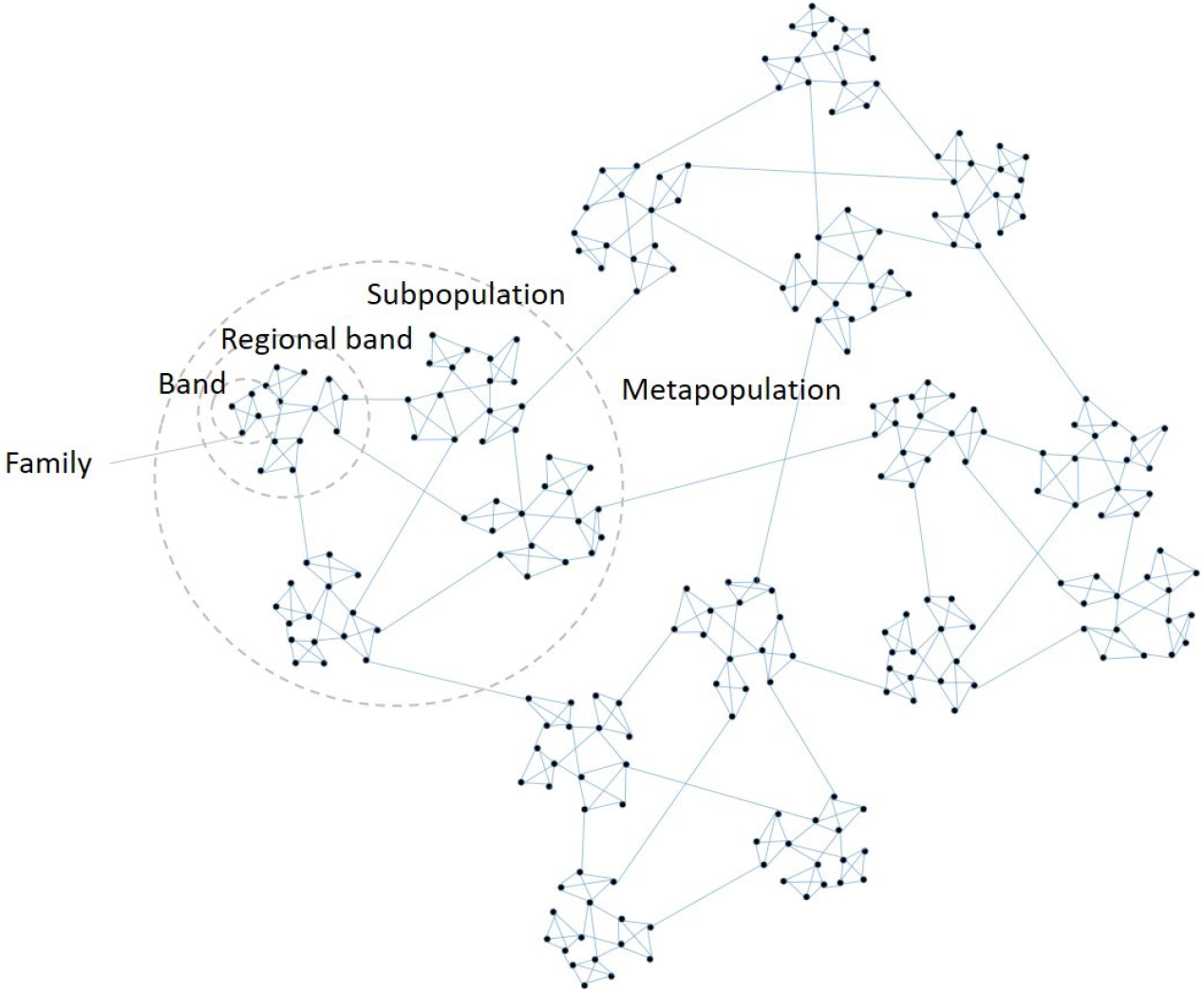
The structure of a hunter-gatherer metapopulation visualized as a hierarchical modular network using the branching statistics from figure 4. Each node in the network is a family (*n* = 256). Four co-residing families form bands (*n* = 64) connected to four other neighboring bands (*n* = 16), which are connected on average to four others forming the sub-populations (*n* = 4), four of which constitute the greater metapopulation (*n* = 1). Fusion-fusion dynamics occur at all levels of the hierarchy as individuals and groups move, interact, aggregate, and disperse at various timescales. Each family exists in a small-world of dense local interactions, but are connected to the larger network by sparse global interactions. As a result, all families are connected to all others by short path lengths that extend across the entire metapopulation.

In hunter-gatherer studies, there is no consensus on the mechanisms that constrain hunter-gatherer group sizes [7, 49, 34, 43]. However, it is useful to consider how the two fundamental currencies of energy and information may come into play in limiting the number of families that co-reside in bands. Consider the finite nature of energy available to human foragers. Data show that hunter-gatherers adjust population *density* (and other adaptive behaviors), rather than population *size* in response to the level of environmental productivity on local landscapes [35, 34, 36] (Figure 4C). This implies that the size of the social group, *g*, (and by extrapolation the entire population structure) is invariant, whereas the size of the home range *H*, is adaptive [36]. As such, the level of intraspecific competition -the number of individuals cooperating and competing -is invariant to local environmental productivity. While competition is invariant, resources are finite ultimately limiting return rates and therefore the ability to maintain the energy demands of foragers and their dependents. Even if we assume return rates, *r*, from cooperative foraging are superlinear, 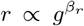, where *β*_*r*_ *>* 1, intraspecific competition in a band will go as *c* ∝ *g*^2^ (equation 9), and so competition outpaces cooperation with increasing group size. The net return is then 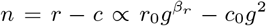, where *β*_*r*_ *>* 1 *<* 2 and *r*_0_ *> c*_0_ (see Figures 5A and B), which the data suggest is maximized at 16 individuals. The upper limit of band sizes is then set by *r* − *c* = 0: in the data the maximum band size is 70 in the Niitsitapi (Blackfoot), an equestrian people of the North American Northern Plains.

As group size increases, not only is there increasing intraspecific competition over energy and resources, but there is also an increasing risk of free riders. To detect free riders an individual must monitor not only their interactions with others, but all interactions in the group, as a free rider may cooperate with some and defect with others. If we assume the cost of monitoring links in the network is constant, *c*_0_, then the per capita cost, *c*, of monitoring group interactions increases quadratically with group size as *c* = *c*_0_*I* = *c*_*I*_*g*^2^. So, like energy, the per capita informational benefits of cooperation will necessarily be less than quadratic, as the exchange of information between individuals cannot be perfect [4, 36]. That is, if there is a constant benefit to interactions among cooperating partners then benefits will increase as, 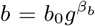, where *β*_*b*_ *<* 2. Therefore, the initial informational benefits to cooperation *b*_0_ *> c*_0_ are quickly outpaced by the costs of policing group interactions. Hunter-gatherer band sizes are thus limited both by competition over energy and information, and these constraints are independent of environmental productivity.

Data suggest hunter-gatherers maximize returns by optimizing local group sizes to ∼ 4 co-residing families. However, by maintaining links with neighboring groups, families exist in a small-world of dense local connections linked to other small-worlds throughout the network by short path lengths; dispersed, low density hunter-gatherer metapopulations are bound by the strength of weak ties [32]. Equation 9 suggests that maintaining the cost of links between groups is much less than within groups as the interaction frequency is far less, in which case the decentralized, modular hierarchical network structure of hunter-gatherer metapopulations maintains connectivity but minimizes the ecological cost of sustaining the total network by minimizing the costs of maintaining links that allow people, material, and information to flow through the network at multiple scales.

## Discussion

This paper demonstrates the central role collective computation plays in structuring hunter-gatherer socioecology. Despite the enormous diversity of hunter-gatherer societies over time and space, cross-cultural ethnographic data show remarkable macroscopic regularities in the scale and structure of hunter-gatherer societies across different environments, reflecting the universal importance of information flow. Hunter-gatherer societies are complex metapopulations comprised of autonomous families who move, fission, and fuse through a nested hierarchy of social groups at multiple levels of social organization at various time scales. These dynamics form a fluid, hierarchically modular social network where individuals, genes, and social information cycle continuously at scales far beyond the local, fine-grained day-to-day interactions of individuals [55, 43, 15]. This complex structure is ultimately shaped by energy constraints on the flow of information among social groups at different levels of social organization resulting in scale-dependent optimizations, modularity, and time-scale separation in dynamics [61].

A key feature of this social structure is the formation of bands – collections of co-residing families bound by norms of reciprocity that constitute the fundamental scale of most daily interactions. From a macroecological and cognitive perspective, bands form collective brains [56] where the information processing of individuals scales up to form distributed cognitive networks at higher levels of aggregation [44, 19]. Band size is a trade-off between the collective benefits of prediction and inference that act to effectively resolve environmental uncertainty versus the ecological maintenance costs of supporting group members, and the policing costs of detecting free-riders.

Data and theory show that the average size of hunter-gatherer bands in the ethnographic record is smaller than predicted for a 60 kg primate; the primate scaling parameters in Table 2 predict hunter-gatherer band sizes of about 32, whereas the empirical average across cultures is 16, half the predicted value [37, 34] (but see [42]). Smaller than predicted social group sizes are compensated by larger than predicted human brain masses such that the collective brain *mass* of a hunter-gatherer band is much as predicted for a 60 kg primate, and is among the the largest of any mammal, regardless of body mass (Figure 3C). Given the tight scaling of the number of cortical neurons and brain mass in both mammals and primates (Figure 1C), hunter-gatherer bands have the highest computational power of any mammal social group. But more importantly, information is exchanged throughout hunter-gatherer metapopulations at scales far beyond the daily interactions of co-residing individuals, and so the information processing capacity of an entire hunter-gatherer metapopulation is clearly unique in the mammalian world. This metapopulation structure thus provides many of the benefits of a large social network (i.e., many potential cooperators; innovators; marriage partners; defenders/raiders; and economies of scale), but minimizes much of the ecological and computational costs of maintaining connectivity.

Hunter-gatherer metapopulations are decentralized networks where the ecological optimizations of energy and information flow that constrain group size play out at local scales. The collective computation of hunter-gatherer societies is to achieve local optima by minimizing band sizes thus minimizing the costs of maintaining network connectivity in a much larger metapopulation. These trade-offs result in small-worlds. The socioecological solution is to maintain large metapopulations by distributing local subpopulations in space. In urban scaling theory, cities are sometimes referred to as *social reactors* [60], a term capturing the hyper-productivity of dense nucleations of people and their interactions concentrated in space. In this spirit, we might then refer to hunter-gatherer societies as *social diffusers*, a term capturing the optimal, adaptive decentralized modularity of people and their interactions dispersed in space. The challenge now is to understand how the social diffusion that characterized the deep evolutionary history of human and hominin social structure over the Plio-Pleistocene eventually nucleated to form the social reactions of the Holocene.

## Conclusions

In the introduction, we posed Tinbergen’s four questions to help frame the adaptive, causative, phylogenetic, and ontogenetic roles of collective computation in hunter-gatherer socioecology. Here, we provide some answers:

### Adaptation

Collective computation is the scale invariant processing of information at multiple spatial-temporal levels of hunter-gatherer social organization. Information is used to build models to make inferences about the world that resolve environmental uncertainty and therefore maximize fitness by optimizing energy budgets, time allocation strategies, reproductive decisions, and survival.

### Causation

Macroscopic regularities are extracted from the environment and used to inform decision-making processes. Optimizations at the individual level scale up to form the collective behavior of social groups integrated into extensive social networks across landscapes to form hierarchically modular metapopulations. Thus, hunter-gatherer societies are not *small-scale* (sensu [6]), but *small-world*.

### Phylogeny

Collective computation in humans has deep phylogenetic roots as the complex 3-dimensional environments of primate evolutionary history promoted large complex brains [62]. The increasingly 2-dimensional niches of the hominin lineage required specialized diets of nutrient-dense resources to fuel the increasingly expensive hominin brain, which in turn required complex predictive models, collective behaviors, and extensive cooperation [14, 48, 39].

### Ontogeny

Information processing at levels above the individual is deeply integrated into all aspects of the human life course, including the scheduling of life history events [48], the social learning of culturally inherited information [8], the mastery of complex skills [74, 52], cooperation amongst non-kin [33, 15], and the coordinated mobility of individuals in space [37, 35].

In hunter-gatherer ecology, collective computation is to information as metabolism is to energy, and both are central to adaptation.

## Acknowledgements

I would like to thank Briggs Buchanan, Hyejin Youn, Chris Kempes, Geoffrey West, Jose Lobo, Eric Rupley, Giovanni Petri, Sam Scarpino, and Rob Walker for invaluable discussions of many of the topics raised in this paper. I would like to thank Fondation IMéRA -Institut d’études avancées, Aix Marseille Université for funding a residential workshop over the summer of 2016 where many of these issues were first discussed. I also thank Tim Kohler, David Wolpert, Darcy Bird, and the Santa Fe Institute for organizing and funding this working group.

